# Structural basis of altered potency and efficacy displayed by a major *in vivo* metabolite of the anti-diabetic PPARγ drug pioglitazone

**DOI:** 10.1101/351346

**Authors:** Sarah A. Mosure, Jinsai Shang, Richard Brust, Jie Zheng, Patrick R. Griffin, Douglas J. Kojetin

**Affiliations:** Skaggs Graduate School of Chemical and Biological Sciences, Scripps Research, Jupiter, Florida 33458, USA; Department of Integrative Structural and Computational Biology, Scripps Research, Jupiter, Florida 33458, USA; Department of Molecular Medicine, Scripps Research, Jupiter, Florida 33458, USA

**Keywords:** ligand-binding protein, drug metabolism, X-ray crystallography, nuclear magnetic resonance (NMR), hydrogen exchange mass spectrometry (HDX-MS), peroxisome proliferator-activated receptor gamma (PPARγ), ligand-binding domain (LBD), nuclear receptor, gene transcription

## Abstract

The thiazolidinedione (TZD) pioglitazone (Pio) is an FDA-approved drug for type 2 diabetes mellitus that binds and activates the nuclear receptor peroxisome proliferator-activated receptor gamma (PPARγ). Although TZDs have potent antidiabetic effects, they also display harmful side effects that have necessitated a better understanding of their mechanisms of action. In particular, little is known about the effect of *in vivo* TZD metabolites on the structure and function of PPARγ. Here, we present a structure-function comparison of Pio and a major *in vivo* metabolite, 1-hydroxypioglitazone (PioOH). PioOH displayed a lower binding affinity and reduced potency in coregulator recruitment assays compared to Pio. To determine the structural basis of these findings, we solved an X-ray crystal structure of PioOH bound to PPARγ ligand-binding domain (LBD) and compared it to a published Pio-bound crystal structure. PioOH exhibited an altered hydrogen bonding network that could underlie its reduced affinity and potency compared to Pio. Solution-state structural analysis using NMR spectroscopy and hydrogen/deuterium exchange mass spectrometry (HDX-MS) analysis revealed that PioOH stabilizes the PPARγ activation function-2 (AF-2) coactivator binding surface better than Pio. In support of AF-2 stabilization, PioOH displayed stabilized coactivator binding in biochemical assays and better transcriptional efficacy (maximal transactivation response) in a cell-based assay that reports on the activity of the PPARγ LBD. These results, which indicate that Pio hydroxylation affects both its potency and efficacy as a PPARγ agonist, contribute to our understanding of PPARγ-binding drug metabolite interactions and may assist in future PPARγ drug design efforts.

---

The thiazolidinedione (TZD) pioglitazone (Pio; brand/trade name Actos) is an FDA-approved drug for the treatment of Type 2 Diabetes Mellitus (T2DM) (1). Pio binds and activates peroxisome proliferator-activated receptor gamma (PPARγ), a nuclear receptor transcription factor that regulates expression of genes important for insulin sensitivity, lipid metabolism, and inflammation (2). PPARγ has the conserved nuclear receptor domain architecture comprised of an N-terminal activation function-1 (AF-1) domain, a DNA-binding domain (DBD), a flexible hinge region, and C-terminal ligand-binding domain (LBD) (3). The DBD recognizes and binds PPAR response elements (PPREs) within the promoter region of target genes, and although DNA binding has been shown to affect receptor function (4–7), the PPARγ LBD has been the primary focus for developing therapeutics because endogenous lipids and synthetic small molecule ligands such as TZDs bind to the LBD and regulate the transcriptional activity of PPARγ.

The PPARγ LBD contains 12 α-helical structural elements that form a three-layer sandwich fold; within this fold is a large hydrophobic core known as the canonical or orthosteric ligand binding pocket. Adjacent to the ligand-binding pocket, a surface formed by the three-dimensional association of helix 3 (h3), helix 4 (h4), helix 5 (h5), and helix 12 (h12) is called the activation function-2 (AF-2) coregulator interaction surface (8,9). Conformational changes in the AF-2 surface that occur in response to ligand binding mediate interactions with coregulators that influence recruitment of chromatin-modifying enzymes and other transcriptional machinery to control downstream gene expression. In the absence of an activating (agonist) ligand, PPARγ recruits transcriptionally repressive corepressor complexes including proteins such as Nuclear Receptor Corepressor 1 (NCoR1) (1,10). However, when an agonist binds to the PPARγ ligand-binding pocket, the AF-2 surface is stabilized, favoring loss of the corepressor complex and recruitment of coactivator proteins such as Thyroid hormone Receptor Associated Protein 220 (TRAP220) (11–13). Previous studies sought to characterize the mechanisms by which agonists activate PPARγ, with early evidence suggesting that h12 acts as a simple “on/off’ switch (14). Recent investigations support more complex mechanisms by which ligands regulate PPARγ function: for instance, by affecting LBD conformational dynamics (11,15,16), binding to an alternate/allosteric site within the LBD (12,17), or regulating post-translational modifications of PPARγ (18,19).

Full PPARγ agonists like the TZDs stabilize an active AF-2 surface conformation through a hydrogen bond network with residues near helix 12 including S289, H323, H449, and Y473 (20). Because Pio is known to have deleterious side effects such as weight gain and musculoskeletal complications, there is demand for a more sophisticated understanding of its effects on PPARγ structure and function (21). One area that requires further investigation is the effects of *in vivo* Pio metabolites. Small molecules like Pio are chemically modified by liver cytochrome P450 (CYP450) enzymes to facilitate excretion (22,23). These modified metabolic intermediates can enter systemic circulation and reach serum levels as high as the original drug (24). Some TZD metabolites maintain the ability to bind and activate PPARγ (23), but enzymatic addition of functional groups may lead to differential regulation of PPARγ activity. Rezulin/troglitazone, another previously FDA-approved TZD for the treatment of T2DM, was removed from the market after a major metabolite caused hepatotoxicity (22). Therefore, it is critical to characterize the metabolites of FDA-approved drugs like Pio as well as the original small molecule.

Previous pharmacokinetic analyses of Pio (Fig. 1A) identified the major *in vivo* metabolite 1-hydroxypioglitazone (PioOH) hyperglycemic effects despite differing from (Fig. 1A), which exhibited weaker anti-Pio only by the addition of a hydroxyl group (25,26). Human studies found that PioOH was present in human serum at equal or greater concentrations than Pio (23), emphasizing its physiological relevance. Here, we investigated how the functional effects of PioOH are related to its structural interaction with PPARγ LBD. Using biochemical and cellular transactivation assays, we showed that Pio hydroxylation reduces potency, as demonstrated by increased EC_50_/IC_50_ values in transactivation and coregulator recruitment assays, which may underlie its reduced anti-hyperglycemic effects relative to Pio *in vivo*. Our crystal structure of PioOH-bound PPARγ LBD identified changes in the ligand-receptor hydrogen bonding network that may underlie its reduced potency, while our solution-state structural analysis using protein NMR and HDX-MS revealed that PioOH stabilized the dynamics of the AF-2 surface compared to Pio. Stabilization of AF-2 surface dynamics were associated with changes in the thermodynamics of coactivator binding and enhanced efficacy in a chimeric cellular transcription assay that reports on the activity of the PPARγ LBD. Overall, these findings contribute to our understanding of how synthetic ligands and their *in vivo* metabolites fine-tune PPARγ function.

**Figure 1.**
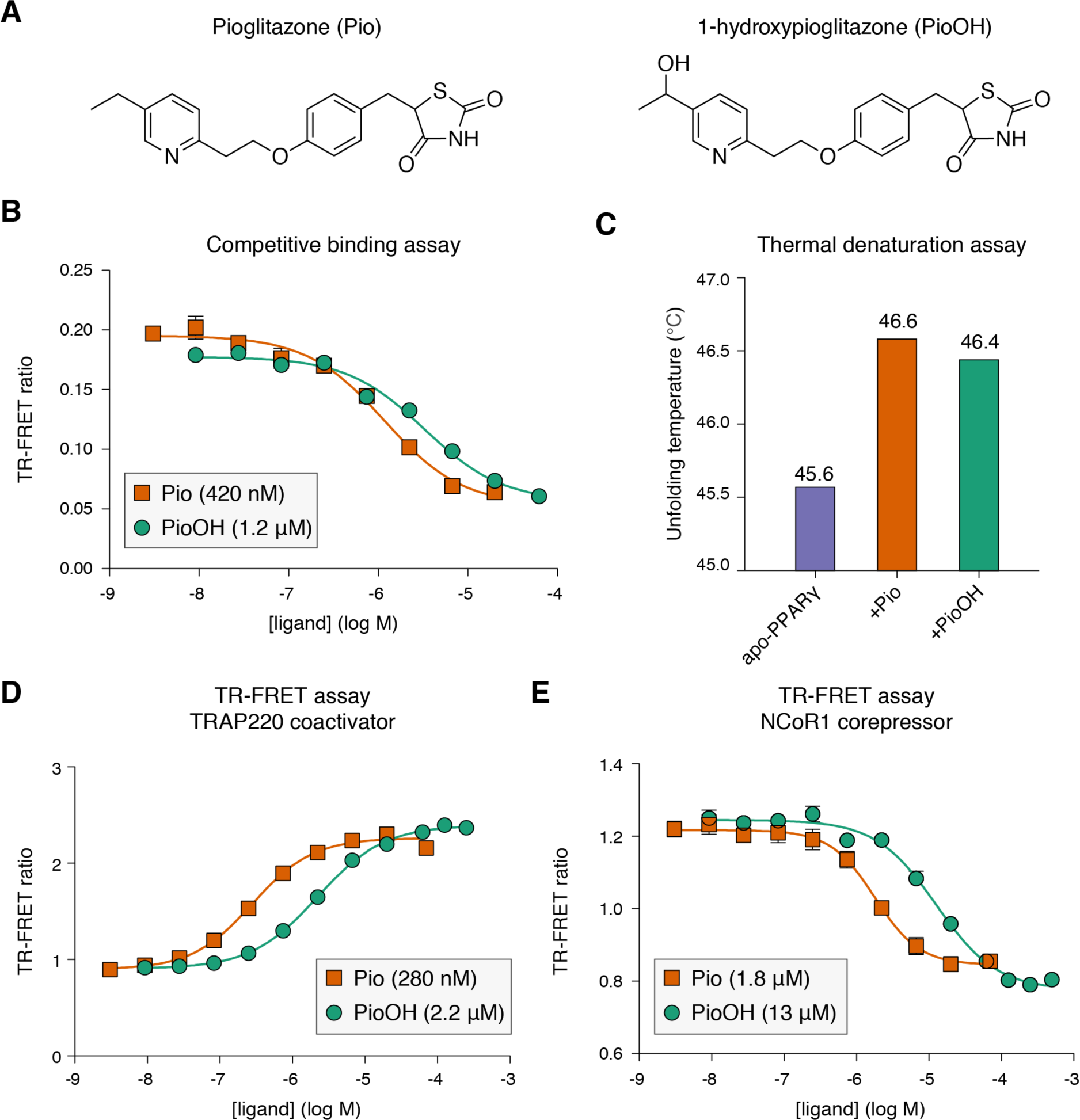
Functional comparison of Pio and PioOH in biochemical assays. (A) Chemical structures of pioglitazone (Pio; left) and the pioglitazone metabolite 1-hydroxypioglitazone (PioOH; right). (B) Competitive binding assay of His-tagged PPARγ LBD with titration of Pio or PioOH. Ligand Ki values for Fluormone™ Pan-PPAR displacement are shown in the legend. (C) T_m_ values from CD spectroscopy thermal denaturation analysis monitored at 222 nm of delipidated apo-PPARγ LBD without or with addition of one molar equivalent of ligand; T_m_ values are noted above the bars. (D) TR-FRET assay of His-tagged PPARγ LBD with FITC-TRAP220 peptide with Pio or PioOH. Ligand EC_50_ values for peptide recruitment are shown in the legend. (E) TR-FRET of His-tagged PPARγ LBD with FITC-NCoR1 peptide titrated with Pio or PioOH. Ligand IC_50_ values for peptide displacement are shown in the legend.

## RESULTS

### PioOH has a weaker binding affinity and coregulator recruitment potency than Pio

We first assessed the binding affinity of Pio and PioOH (Fig. 1A) to the PPARγ LBD using a fluorescent ligand displacement assay (Fig. 1B), which revealed that the metabolic conversion of Pio to PioOH via addition of a hydroxyl group to the end of the hydrophobic side-chain of Pio weakened the interaction with the PPARγ LBD (P<0.001). Circular dichroism (CD) spectroscopy thermal denaturation experiments confirmed binding and revealed that the unfolding temperature of the PPARγ LBD is increased upon binding both Pio and PioOH (Fig. 1C), indicating binding of these ligands stabilized the LBD. We also performed time resolved fluorescence resonance energy transfer (TR-FRET) biochemical assays to determine how Pio and PioOH affect the interaction with peptides derived from a PPARγ-interacting transcriptional coactivator (TRAP220) and corepressor (NCoR1). Consistent with its reduced binding affinity, PioOH was less potent than Pio in recruiting the TRAP220 peptide (Fig. 1D) and displacing the NCoR1 peptide (P<0.001) (Fig. 1E).

### Crystal structure of PioOH-bound PPAR*γ* LBD reveals an altered hydrogen bonding network

To determine the structural basis of PioOH activity compared to Pio, we solved the crystal structure of PPARγ LBD bound to PioOH at 1.88 Åresolution using molecular replacement (Table 1). The X-ray crystal structure of Pio bound to PPARγ LBD was recently solved at comparable resolution (1.801 Å; PDB code 5Y2O) (20). Both Pio and PioOH-bound PPARγ LBDs crystallized as dimers: h12 in chain Λ adopts an “active” conformation with the ligand bound in the orthosteric pocket, whereas h12 in chain B adopts an atypical conformation in which h12 docks onto adjacent molecules within the crystal lattice. Structural alignment of chain Λ in the Pio and PioOH structures resulted in an overall RMSD of 0.385 Å (Fig. 2A), indicating there were no overall structural changes. However, subtle structural changes (Fig. 2B) that were likely influenced by differences in the ligand-receptor hydrogen bond network (Fig. 2C,D) could explain the reduced affinity of PioOH.

**Figure 2.**
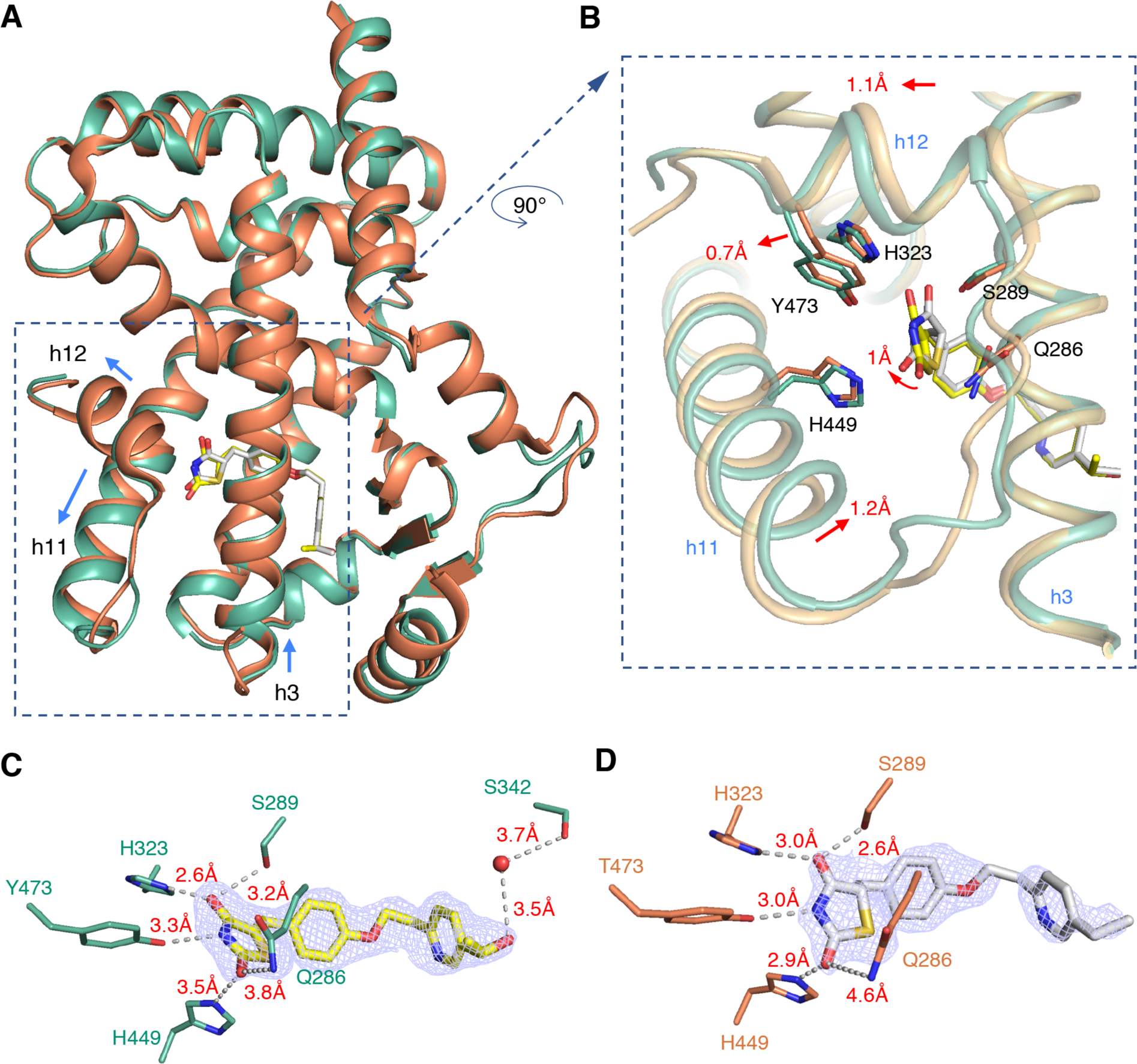
Comparison of PPARγ LBD crystal structures bound to Pio or PioOH. (A) Structural alignment of PioOH-bound PPARγ LBD crystal structure (PDB code 6DHA, chain A; green cartoon, yellow ligand) and Pio-bound PPARγ LBD crystal structure (PDB code 5Y2O, chain A; orange cartoon, white ligand). (B) 90° counter-clockwise rotation and zoomed in view of the region of the ligand-binding pocket that contacts the Pio and PioOH TZD head groups. The side chains of residues that form hydrogen bonds with TZD head group are shown as sticks, and red arrows indicate structural shifts in secondary structure between the Pio and PioOH-bound structures. (C,D) Omit maps (2FO-FC, contoured at 1 σ) of (C) PioOH and (D) Pio displayed with hydrogen bonds to residues within the ligand-binding pocket (gray dotted line and red text), as well as a water molecule (red sphere) in the PioOH structure that participates in a water-mediated hydrogen bond between the S342 side chain and the PioOH hydroxyl group (D).

**Table 1.** Crystallography refinement statistics.

| | PPARγ LBD bound to 1-hydroxypioglitazone (PioOH) |
| --- | --- |
| Data collection | BCSB-5.0.2 |
| Space group | C 1 2 1 |
| Cell dimensions |  |
| $a, b, c$ (Å) | 93.08, 61.85, 119.67 |
| $\alpha, \beta, \gamma$ (°) | 90, 102.73, 90 |
| Resolution | 49.12–1.88 (1.95–1.88) |
| $R_{\text{pim}}$ | 0.056 (0.456) |
| $I / \sigma(I)$ | 4.93 (1.29) |
| $CC1/2$ in highest shell | 0.618 |
| Completeness (%) | 97.33 (95.12) |
| Redundancy | 1.8 |
| Refinement |  |
| Resolution (Å) | 1.88 |
| No. of reflections | 98149 |
| $R_{\text{work}}/R_{\text{free}}$ (%) | 22.05/25.53 |
| No. of atoms |  |
| Protein | 4147 |
| Water | 250 |
| $B$-factors | |
| Protein | 46.19 |
| Ligand | 52.84 |
| Water | 45.78 |
| Root mean square deviations |  |
| Bond lengths (Å) | 0.008 |
| Bond angles (°) | 0.95 |
| Ramachandran favored (%) | 96.08 |
| Ramachandran outliers (%) | 1.37 |
| PDB accession code | 6DHA |
| *Values in parentheses indicate highest resolution shell.* |  |

The TZD head group of both ligands form hydrogen bonds with the side chains of residues S289, H323, H449, and Y473. The hydrogen bond lengths in the PioOH-bound structure (Fig. 2C) were extended relative to the Pio-bound structure (Fig. 2D) for S289, H323, and Y473. The PioOH TZD head group was tilted ~1 Å such that these interacting side chains were farther from their hydrogen bond donor/acceptor, while Q286 and H323 were closer in the PioOH-bound structure despite overall minimal changes in side chain conformations. Additionally, the PioOH hydroxyl group formed a weak water-mediated hydrogen bond with the side chain of S342 (Fig. 2C), which likely contributed to the change in binding mode that shifted the TZD head group of PioOH relative to Pio. The PioOH TZD head group displacement was reciprocated by a 1.1 Å shift of h12 away from h3 and a 1.2 Å shift of h11 toward h3 (Fig. 2A,B). The introduction of the hydroxyl group in PioOH shifted the binding pose of the terminal methyl group towards G284 on helix 3. However, this methyl shift did not introduce any steric clashes with residues near helix 3. Thus, it is possible that the combination of the weakened PioOH hydrogen bond interactions with residues near helix 12, as well as the weak water-mediated hydrogen bond introduced by the hydroxylation of Pio, contributes to the weakened affinity and potency of PioOH.

### Solution structural analysis reveals enhanced AF-2 stabilization by PioOH

Although ligand-bound crystal structures provide important molecular detail into the binding mode of PPARγ ligands, it has been difficult to identify structural mechanisms of graded ligand activity, such as full vs. partial agonism, due to the structural homogeneity of protein conformations within the crystals (27). Typically, backbone conformations observed in ligand-bound PPARγ crystal structures are highly similar due to crystal packing forces that favor the most stable crystallized conformation, and thus they are not representative of the ensemble of conformations present in solution. Alternatively, structural studies that probe the dynamics of proteins in solution, such as NMR spectroscopy and HDX-MS, have identified more nuanced yet functionally relevant characteristics of graded activation (11,15,16).

We performed differential NMR analysis by collecting 2D [^1^H,^15^N]-TROSY-HSQC NMR spectra of ^15^N-labeled PPARγ LBD with Pio or PioOH (Fig. 3A). NMR peaks were assigned to residues using the minimal NMR chemical shift method (28) based on their nearest neighbor to rosiglitazone-bound PPARγ LBD NMR chemical shift assignments (11). This method allowed us to assign 181 well-resolved peaks for PioOH and Pio; 93 residues remained unassigned mostly due to NMR peak overlap that made transfer of NMR assignments difficult (Fig. 3B). We calculated NMR chemical shift perturbations (CSPs) between the Pio- and PioOH-bound states to identify changes in the chemical environment of PPARγ LBD (Fig. 3C), which can be due to ligands as well as allosteric conformational differences in residues that directly contact the differences that occur between Pio- and PioOH-bound PPARγ LBD. There were eight residues with CSPs more than two standard deviations (CSP > 0.037 p.p.m.) from the mean CSP (0.013 p.p.m.) (Fig. 3B,C). These residues were located primarily in the β-sheet region (L340, I341, G346, and V248) (Fig. 4A and B), or at the putative site of ligand entry/exit (29), a region comprised of the h6-7 loop (F360 and D362) and the N-terminus of h3 (A278 and I279) (Fig. 4C and D). Another residue with a significant CSP (I262) was located in the Ω-loop, a flexible region between h2 and h3 absent from most PPARγ LBD crystal structures.

**Figure 3.**
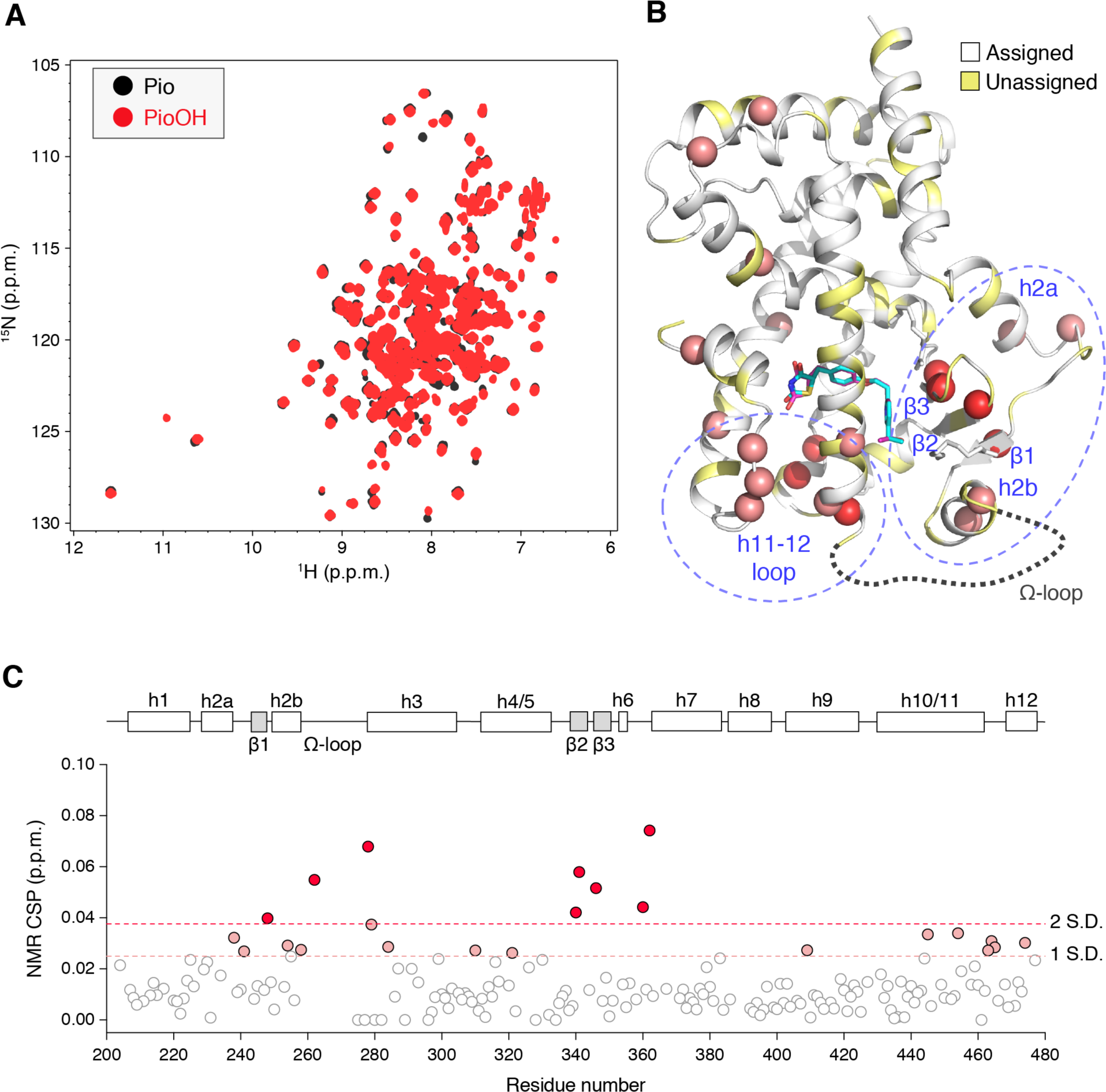
Differential NMR analysis of Pio- and PioOH-bound PPARγ LBD. (A) Overlay of 2D [^1^H,^15^N]-TROSY-HSQC spectra ^15^N-labeled PPARγ LBD (200 μM) with 2 molar equivalents of Pio or PioOH. (B) Residues with NMR chemical shift perturbations (CSPs) greater than 1 S.D. from the average plotted on the PPARγ LBD with the Pio and PioOH ligands displayed as blue and magenta sticks, respectively. Blue dashed ovals indicate two regions with the most CSPs, the β-sheet region and the putative pocket entry/exit region. The dashed gray loop indicates the conformationally flexible Ω-loop absent from in the crystal structures. Pio is shown in light blue and PioOH is shown in magenta. (C) NMR CSPs plotted by residue; residues are highlighted with CSPs > 1 S.D. (pink dashed lines and circles) or 2 S.D. (red dashed line and circles) from the mean CSP (0.013 p.p.m.). PPARγ LBD structural elements are depicted linearly above the graph.

**Figure 4.**
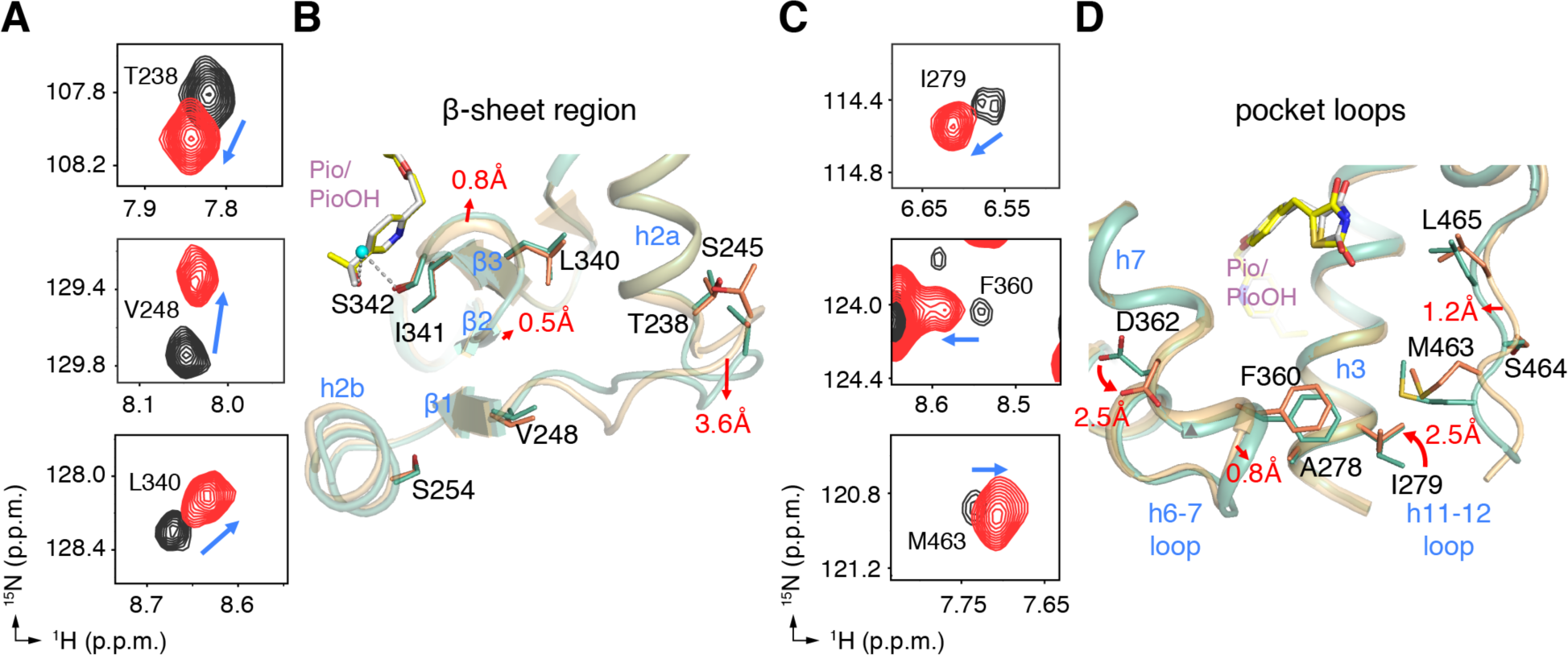
Correlation between differential NMR data and crystal structures. (A) Representative residues with notable NMR CSPs in the β-sheet region of the ligand-binding pocket. Blue arrows indicate the direction of the CSP from Pio (black peaks) to PioOH (red peaks). (B) Subtle conformational changes in β-strand region are observed in PPARγ LBD crystal structures bound to Pio (PDB code 5Y2O, chain A; light orange cartoon, white ligand) and PioOH (PDB code 6DHA, chain A; light green cartoon, yellow ligand). (C) Representative residues, depicted as in (A), with notable CSPs in the ligand entry/exit region of the ligand-binding pocket, which (D) also manifest as subtle conformational changes in ligand-bound crystal structures, as depicted in (B).

The CSPs between Pio and PioOH-bound PPARγ observed for the β-sheet region were consistent with the water-mediated hydrogen bond observed in the PioOH-bound crystal structure between the PioOH hydroxyl group and the Ser342 side chain, which is not present in the Pio-bound structure. Although the crystal structures only showed minor structural changes (0.5-0.8 Å) in the backbone and negligible side chain alterations, the water-mediated hydrogen bond showed a pronounced effect on the chemical environment of residues this region as detected by NMR (Fig. 4A,B and 3B). At the same time, residues with CSPs near the h6-7 loop were reflected by structural changes in the crystal structures: there was a 0.8 Åshift of the h6-7 loop toward h3 accompanied by a downward shift of similar magnitude for the F360 side chain, and the side chains of D362 and I279 side chains showed a 2.5 Åshift (Fig. 4D). Together, the CSP data suggest that hydroxylation of Pio has the greatest effect on the conformation of the ligand entry/exit site and the β-strand region of the PPARγ LBD, but only some of these changes were apparent in the crystal structure.

In addition to providing information about conformational changes via CSP analysis, NMR spectra can offer insight into differences in protein dynamics between two ligand-bound forms. For example, in our differential NMR analysis, a change in NMR lineshape, or peak intensity, indicates a change in dynamics on the microsecond-millisecond (μs-ms) timescale (i.e., intermediate exchange on the NMR timescale between two or more conformations). A number of residues in PioOH-bound PPARγ LBD showed an increase in NMR peak intensity compared to Pio-bound PPARγ LBD, indicating PioOH stabilized μs-ms timescale dynamics introduced by Pio (Fig. 5A-C). Importantly, many of the stabilized residues were located at the AF-2 coactivator binding surface, indicating that PioOH may more effectively stabilize h12 docking against h3 and h11 in the AF-2 “active” conformation that is compatible with coactivator binding.

**Figure 5.**
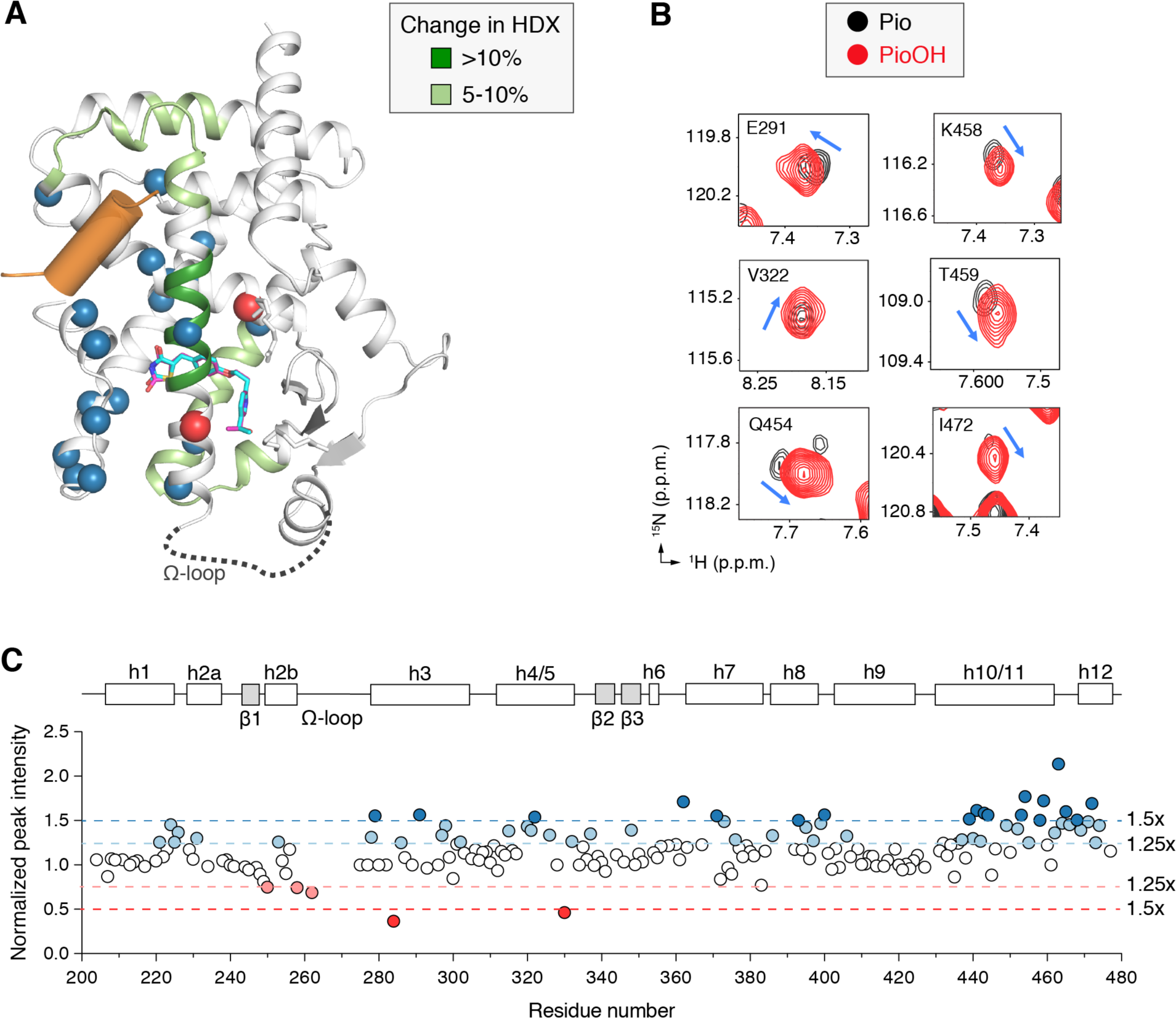
PioOH enhances stabilization of the AF-2 coregulator interaction surface. (A) Results of HDX-MS and NMR peak intensity analysis displayed on PPARγ LBD crystal structure with a coactivator peptide (orange cylinder) bound to the AF-2 surface (PDB code 2PRG, chain A); Pio and PioOH ligand binding modes are displayed as blue and magenta sticks, respectively. Regions of the PPARγ LBD that exhibit increased protection from HDX bound to PioOH relative to Pio are colored light or dark green on the cartoon diagram. Residues with increased (blue spheres) or decreased (red spheres) NMR peak intensity when PPARγ LBD is bound to PioOH, relative to PPARγ LBD bound to Pio, in the differential NMR analysis are also displayed. (B) Representative residues with increased NMR peak intensity in the PioOH-bound form relative to the Pio-bound form. (C) Differential NMR peak intensity analysis plotted by residue. PioOH-bound peak intensities are normalized to Pio-bound peak intensities; residues are highlighted with normalized peak intensities greater than (blue) or less than (red) 1.25x and 1.5x the average normalized peak intensity difference. PPARγ LBD structural elements are depicted linearly above the graph.

Whereas NMR lineshape analysis reports on μs-ms timescale dynamics, HDX-MS probes molecular “breathing” motions or dynamics on a timescale of seconds (30). To determine whether the enhanced AF-2 stabilization observed for PioOH by NMR can be detected at longer timescales, we performed differential HDX-MS of PPARγ LBD bound to Pio or PioOH (Fig. 5A; Table S1). Consistent with our NMR analysis, PioOH afforded greater protection from deuterium exchange than Pio for peptides within h3 and the h3-h4/5 loop within the AF-2 surface. There was also stabilization of the h6-7 loop region, which exhibited significant NMR CSPs and stabilization of μs-ms time scale dynamics when bound to PioOH, inferred by increased NMR peak intensities.

### PioOH differentially affects coregulator binding mechanisms

Although our binding assays showed that PioOH is less potent than Pio, our NMR and HDX-MS analyses revealed that PioOH more effectively stabilizes the AF-2 surface. AF-2 stabilization is linked to enhanced functional agonism through higher affinity coactivator binding (31). To test this directly, we performed fluorescence polarization (FP) assays to determine how Pio hydroxylation affects the binding affinity of PPARγ LBD for the TRAP220 coactivator peptide. Compared to apo-PPARγ LBD, both ligands increased the affinity of the TRAP220 peptide relative to apo-PPARγ LBD. PioOH-bound PPARγ LBD consistently bound to the TRAP220 coactivator peptide with higher affinity than Pio-bound PPARγ LBD in each trial, but the difference in the fitted K_d_ values did not achieve statistical significance (P~0.1) (Fig. 6A).

**Figure 6.**
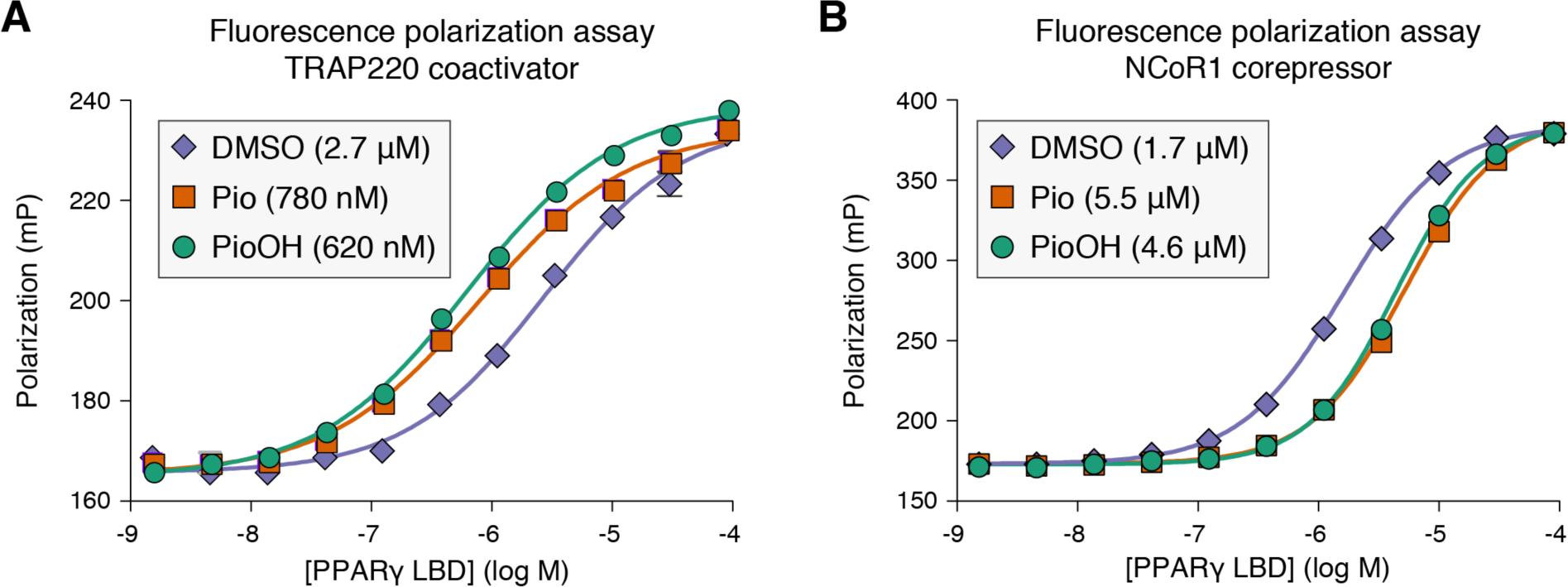
Pio hydroxylation affects coactivator and corepressor binding to PPARγ LBD. (A,B) Fluorescence polarization assays determined the binding affinities of FITC-labeled peptides derived from the (A) TRAP220 coactivator and (B) NCoR corepressor. Assays performed using a saturating amount of ligand, or vehicle control.

We also tested the effect of PioOH vs. Pio on corepressor binding affinity. Using FP assays with a FITC-labeled NCoR1 corepressor peptide, we found that both ligands reduced the affinity of the NCoR1 peptide relative to apo-PPARγ LBD (Fig. 6B). However, in this case PioOH was significantly less effective than Pio at reducing NCoR binding (P<0.001). Taken together, this indicates that although the ligand-dependent effects on coactivator and corepressor binding affinities may be related, where TZD agonists both strengthen TRAP220 coactivator affinity and weaken NCoR1 corepressor affinity relative to apo-PPARγ, they are not directly correlated since the ligand that produced marginally improved TRAP220 coactivator affinity (PioOH) did not weaken NCoR1 corepressor affinity the most.

Although the FP assays did not reveal a significantly enhanced coactivator binding affinity, the polarization window for TRAP220 binding produced by PioOH was consistently greater than that of Pio (P<0.05), suggesting a greater reduction in molecular tumbling or better stabilization of peptide binding, perhaps via better AF-2 stabilization, at saturating conditions. To test the thermodynamic basis of this observation, we performed isothermal titration calorimetry using the unlabeled TRAP220 peptide and the PPARγ LBD bound to two molar equivalents of either Pio or PioOH (Fig. 7A and B). Consistent with the FP assay, the K_d_ of peptide binding was lower for PioOH-bound PPARγ, but did not achieve statistical significance; however, the enthalpy (ΔH) of binding was significantly reduced compared to Pio, suggesting stabilized electrostatic contacts with TRAP220 when bound to PioOH (Table 2 and Fig. 7C). Additionally, the entropy (ΔS) of binding was more negative and thus less favorable for PioOH, indicating that the binding dynamics between TRAP220 and PioOH-bound PPARγ LBD are less flexible than for the Pio-bound structure.

**Figure 7.**
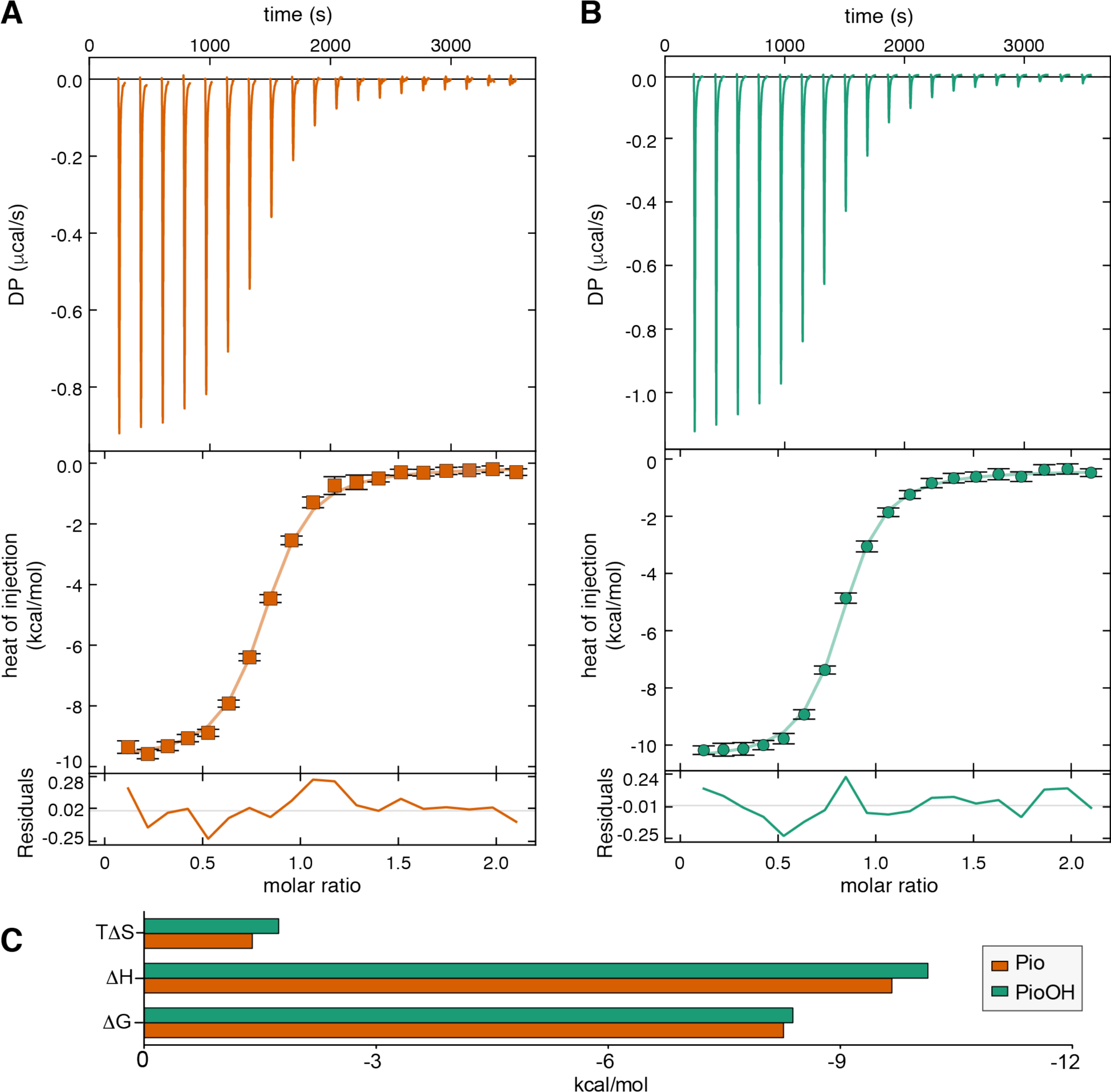
PioOH stabilizes coactivator binding to the PPARg LBD. (A,B) Representative thermograms and normalized plotted data from ITC analysis of TRAP220 binding to Pio-(A) or PioOH-(B) bound PPARγ LBD. (C) Fitted thermodynamic parameters from a global analysis of two replicate runs per condition.

**Table 2.** Fitted isothermal titration calorimetry parameters for TRAP220 titrated into PPARγ LBD.

| Ligand | K<sub>d</sub> (nM) | $\Delta G$ (kcal/mol) | $\Delta H$ (kcal/mol) | $\Delta S$ (kcal/mol*K) |
| --- | --- | --- | --- | --- |
| Pio | 871<br>(68.3% CI: 740 to 1020) | -8.39 | -10.136<br>(68.3% CI: -10.3589 to -9.9186) | -5.856 |
| PioOH | 708<br>(68.3% CI: 606 to 823) | -8.27 | -9.668<br>(68.3% CI: -9.8940 to 9.4541) | -4.669 |
| 68.3% CI = confidence interval provided by SEDPHAT analysis |  |  |  |  |

### PioOH enhances the transcriptional activity of the PPARγLBD

To compare the cellular activation properties of Pio and PioOH, we performed transcriptional reporter assays by transfecting HEK293T cells with an expression plasmid encoding full-length PPARγ along with a reporter plasmid containing three copies of the PPAR DNA response element (3xPPRE) upstream of the luciferase gene. Both Pio and PioOH caused a concentration-dependent increase in PPARγ transcription, and consistent with the biochemical binding data PioOH showed reduced cellular potency relative to Pio (P<0.0001) (Fig. 8A).

**Figure 8.**
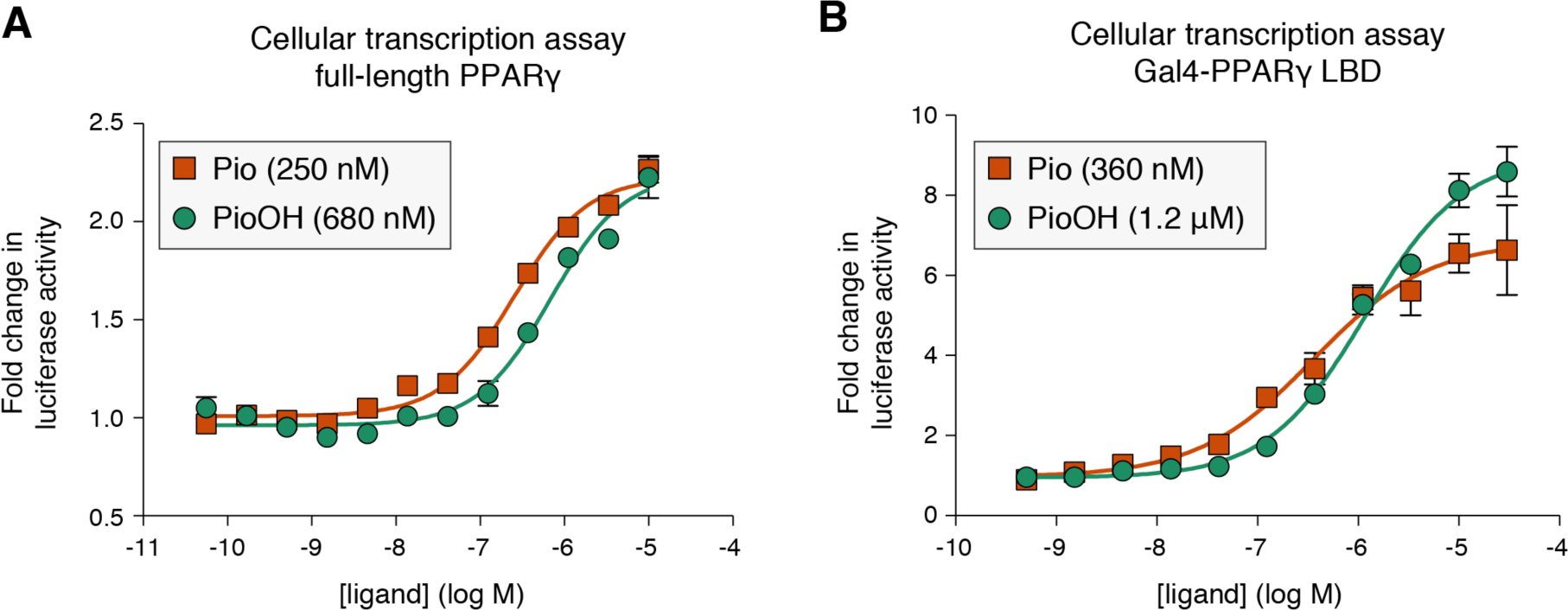
PioOH induces a modestly greater transcriptional efficacy of the PPARγLBD. (A) Full-length PPARγ luciferase transcriptional assay using a 3xPPRE-luciferase reporter plasmid in HEK293T cells treated with increasing concentrations of Pio or PioOH; data are normalized to DMSO control treated cells. (B) PPARγ LBD-Gal4 DBD luciferase transcriptional assay using a 5xUAS-luciferase reporter plasmid in HEK293T cells treated with increasing concentrations of Pio or PioOH; data are normalized to DMSO control treated cells. Ligand EC_50_ values for cellular transcriptional activation are shown in the legends.

We also performed a cell-based transcriptional reporter assay using a chimeric expression construct comprised of the PPARγ LBD fused to the Gal4 yeast transcription factor DBD. This construct lacks the N-terminal ligand-independent activation function-1 (AF-1) and native DNA-binding domain in PPARγ and thus reports on LBD activity only, and provides a higher window of activation and is therefore more sensitive to graded levels of agonism (11) compared to the full-length PPARγ assay. We transfected HEK293T cells with the Gal4-PPARγ LBD expression plasmid and a reporter plasmid containing five tandem repeats of the Gal4 Upstream Activation Sequence (5xUAS) placed upstream of the luciferase gene. Consistent with the full-length PPARγ results, PioOH was less potent than Pio (P<0.02) (Fig. 8B). Intriguingly, however, PioOH yielded a significantly greater maximal luciferase activity than Pio (P<0.05). Thus, enhanced AF-2 surface stabilization afforded by PioOH corresponds to increased transcriptional activition of the LBD-only construct, which may not be apparent in the full-length assay due to lower sensitivity or contributions of the N-terminal ligand-independent AF-1 domain.

## DISCUSSION

Pioglitazone (brand/trade name Actos) remains an important option for treatment of T2DM because of its potent insulin sensitizing effects (32,33). However, adverse side effects have warranted further investigation into its mechanism of action. To our knowledge, little is known how Pio metabolites, or the metabolites of any other PPARγ-binding drug, affect the structure and function of PPARγ. Therefore, we performed a comparative structure-function analysis of Pio and its major *in vivo* Pio metabolite, 1-hydroxypioglitazone (PioOH), using X-ray crystallography and solution-state structural methods in combination with biochemical and cell-based assays. We found that PioOH binds PPARγ with weaker affinity than Pio, but produces a modestly more efficacious agonist response at saturating ligand conditions in biochemical (TR-FRET) and cellular (chimeric Gal4 transactivation) assays, potentially due to stabilization of an LBD conformation with enhanced coactivator binding affinity. Crystal structures of PPARγ LBD bound to Pio and PioOH did not show any overall structural changes that could lead to the improved agonist profile of PioOH. However, solution-state structural analysis using NMR spectroscopy and HDX-MS revealed that PioOH better stabilizes the dynamics of the AF-2 surface, which is consistent with its improved biochemical, thermodynamic, and cellular agonist features over Pio on LBD activity. Because Pio and PioOH concentrations are similar in human serum (23), the reduced potency of PioOH likely underlies its reduced anti-hyperglycemic efficacy relative to Pio (25,26). If PioOH serum concentrations were higher than Pio, it is possible that PioOH would be nearly as efficacious as Pio.

Analysis of the metabolites of other nuclear receptor ligands has provided essential insight to into the activity of the parent compound/drug and informed the design of small molecule therapeutics. For example, testosterone is the primary endogenous ligand of androgen receptor (AR), but in tissues expressing 5α-reductase, testosterone is converted to 5α-dihydrotestosterone (5α-DHT), which binds with greater selectivity and affinity to AR and is thought to enhance androgenic effects in specific tissues such as the prostate and seminal vesicles (34). Intriguingly, 5α-DHT differs from testosterone only by the presence of an additional unsaturated double bond and, similar to our analysis here of PPARγ, crystal structures of AR LBD bound to 5α-DHT or testosterone revealed no large structural changes (overall Cα RMSD only 0.238 Å) that would lead to improved function (35). However, comprehensive analysis of atomic geometries within the AR ligand-binding pocket revealed the subtle interactions in the two liganded complexes that determine AR ligand affinity. These findings and others informed the development of selective androgen receptor modulators (SARMs) with more favorable ratios of anabolic to androgenic effects (36,37)

Drug metabolite studies have also informed the development of selective estrogen receptor modulators (SERMs). Pharmacokinetic analysis revealed that the metabolites of tamoxifen had variable activity at Estrogen Receptor (ER): 4-hydroxytamoxifen and N-desmethyl-4-hydroxytamoxifen (endoxifen) exhibited 100-fold greater affinity and were found to underlie tamoxifen’s ability to antagonize ER in breast cancer tissue (38,39). Another metabolite of tamoxifen was found to have weak antiestrogenic activity, and equivalent chemical modifications were made to the SERM Toremifene to produce Ospemifene, a SERM with promising anti-tumor and pro-bone growth effects (40). Ultimately, structural studies probing the molecular and biophysical basis of ER-metabolite interactions have contributed to SERM identification and characterization (39,41).

Efforts to develop selective PPAR modulators (SPPARMs) have been underway for over a decade (42). Recent work has focused on separating the “classical” agonist effects of PPARγ-binding ligands from their ability to regulate “non-classical” posttranslational modifications of PPARγ (19,43). Here, our work indicates that the introduction of a water-mediated hydrogen bonding network underlies the weakened binding affinity of PioOH relative to Pio. This was somewhat unexpected because the hydrogen bonding ability introduced by the addition of the hydroxyl group in PioOH in an otherwise hydrophobic ligand side-chain (in Pio) would typically be considered as overall favorable for binding affinity, but underlies the notion that hydrophobic effects also play an important role in ligand binding affinity (44). However, although PioOH displayed weaker binding affinity to Pio, our solution-based structural analysis revealed that PioOH better stabilized several regions of the LBD than Pio. The stabilized regions include the AF-2 coregulator surface and a region of helix 7 that includes a ligand-dependent SUMOylation site (Lys367) implicated in promoting PPARγ-mediated repression of pro-inflammatory genes (45). The water-mediated hydrogen bond observed between PioOH and S342 side-chain in our crystal structure was confirmed by our NMR studies, which showed conformational changes via chemical shift changes for residues within the β-sheet region of the ligand-binding pocket. Ligand interactions with the backbone amide of Ser342 have previously been implicated in inhibiting the phosphorylation of S273 (S245 in PPARγ isoform 1), which is associated with obesogenic side effects of PPARγ activation (18,19). Thus, it will be interesting to determine in future studies if the Pio metabolite, PioOH, enhances the non-classical effects of Pio on PPARγ activity. Taken together, our findings provide new insight into how structure-function studies of ΡΡΑΚγ-binding drug metabolites may help to inform on the function and design of PPARγ ligands.

## EXPERMIENTAL PROCEDURES

### Ligands and protein preparation

Pioglitazone (Pio; Cayman Chemical) and 1-hydroxypioglitazone (PioOH; Axon Medchem) were prepared in DMSO-d6 as 50 mM and 7 mM stocks, respectively. Human PPARγ LBD (residues 203–477, isoform 1 numbering) was expressed in *Escherichia coli* BL21(DE3) cells as TEV-cleavable hexahistidine (6xHis)-tagged fusion protein using protocols previously described (11,12). Purified protein was delipidated using LIPIDEX 1000 resin (Perkin Elmer) and stored in 20 mM potassium phosphate (pH 7.4), 50 mM potassium chloride, 0.5 mM EDTA, and 5 mM TCEP. FITC-labeled peptides derived from the TRAP220 coactivator (residues 638–656; NTKNHPMLMNLLKDNPAQD) and NCoR1 corepressor (residues 2256–2278; DPASNLGLEDIIRKALMGSFDDK) were synthesized by Lifetein; peptides contained a six-carbon linker (Ahx) after the FITC label, and the C terminus was amidated for stability.

### Circular dichroism (CD) spectroscopy

CD wavelength scans and thermal denaturation experiments monitored at 222 nm were performed in CD buffer (10 mM potassium phosphate (pH 7.4) and 50 mM potassium fluoride) to determine the folding and stability of PPARγ LBD (10 μM) in the presence of one molar equivalent of Pio or PioOH on a Jasco J-815 spectropolarimeter.

### TR-FRET competitive ligand displacement and coregulator interaction assays

Time-resolved fluorescence resonance energy transfer (TR-FRET) assays were performed in black low-volume 384-well plate (Greiner) using a buffer containing 20 mM potassium phosphate (pH 7.4), 50 mM potassium chloride, 0.5 mM EDTA, and 5 mM TCEP, and 0.01% Tween-20. For the ligand displacement assay, each well (22.5 μL per well) contained 1 nM 6xHis-PPARγ LBD protein, 1 nM LanthaScreen Elite Tb-anti-His Antibody (Thermo Fisher Scientific), and 5 nM Fluormone Pan-PPAR Green tracer ligand (Invitrogen) in TR-FRET buffer. For the TR-FRET coregulator interaction assay, each well contained 400 nM FITC-labeled TRAP220 or NCoR1 peptides, 4 nM 6xHis-PPARγ LBD protein, 1 nM LanthaScreen Elite Tb-anti-His Antibody (Thermo Fisher Scientific), and 400 nM peptide in TR-FRET buffer in 22.5 μL total volume per well. Ligand stocks were prepared via serial dilution in DMSO, added to wells in triplicate to a final DMSO concentration of 1%, and the plates were incubated at 25 °C for ~1 h and read using BioTek Synergy Neo multimode plate reader. The Tb donor was excited at 340 nm; its fluorescence emission was monitored at 495 nm, and the acceptor FITC emission was measured at 520 nm. The TR-FRET ratio was calculated as the signal at 520 nm divided by the signal at 495 nm. Data were plotted using GraphPad Prism and fit to the appropriate equation: for the ligand displacement assay, data were fit to a competitive one site fit K_i_ equation using the known binding affinity of Fluormone^TM^ Pan-PPAR Green tracer ligand (2.8 nM; Thermo Fisher Scientific product insert PV4894); and for the coregulator interaction assay data were fit to a sigmoidal dose response equation.

### Cell-based transactivation assays

HEK293T cells (ATCC; cat# CRL-3216) cultured in DMEM media (Gibco) supplemented with 10% fetal bovine serum (FBS) and 50 units mL^-1^ of penicillin, streptomycin, and glutamine were grown to 90% confluency in a T-75 flask before seeding 4 million cells per well in 10-cm dishes. Seeded cells were transfected using transfection reagent containing 27 μL X-treme Gene 9 (Roche) in serum-free Opti-mem reduced serum media (Gibco) with either 4.5 μgpCMV6-XL4 plasmid containing full-length human PPARγ2 and 4.5 μg 3X multimerized PPRE-luciferase reporter or 4.5 μg Gal4-PPARγ LBD and 4.5 μg 5X Upstream Activation Sequence (UAS) luciferase reporter. After 18 hrs incubation at 37 °C in a 5% CO2 incubator, the transfected cells were plated in quadruplicate in white 384-well plates (Perkin Elmer) at a density of 10,000 cells per well (20 μL volume) and incubated 4 hrs then treated with 20 μL of vehicle control (1% DMSO in DMEM media) or 1:2 serial dilution of each compound from 56 pM-10 μΜ (1% final DMSO concentration). After 18 hrs, luciferase activity was measured by addition of 20 μL Britelite Plus (Perkin Elmer) and luminescence was read using a BioTek Synergy Neo multimode plate reader. Data were plotted in GraphPad Prism as fold change in luminescence of compound-treated cells over DMSO-treated control cells vs ligand concentration and fit to a sigmoidal dose response equation.

### Fluorescence polarization coregulator interaction assays

Fluorescence polarization assays were performed in black low-volume 384-well plates (Greiner) using a buffer containing 20 mM potassium phosphate (pH 7.4), 50 mM potassium chloride, 0.5 mM EDTA, 5 mM TCEP, and 0.01% Tween-20. Each well contained 100 nM FITC-labeled TRAP220 coactivator peptide (Lifetein), a serial dilution of PPARγ LBD (1.5 nM-90 μM), with a fixed concentration of vehicle control (1% DMSO) or ligand equal to the highest protein concentration (90 μMPio or PioOH) in triplicate. Plates were incubated 2 hrs at 4°C and read using BioTek Synergy Neo multimode plate reader. Data were plotted in GraphPad Prism and fit to a sigmoidal dose response equation.

### Statistical Tests

Statistical significance between best fit EC_50_, IC_50_, and top values in biochemical and cell-based assays was determined using the GraphPad Prism Extra sum-of-squares F test analysis with alpha equal to 0.05.

### Isothermal titration calorimetry

A peptide derived from the TRAP220 coactivator (residues 638-656; NTKNHPMLMNLLKDNPAQD) was synthesized by Lifetein and resuspended at 500 μM in buffer containing 20 mM potassium phosphate (pH 7.4), 50 mM potassium chloride, 0.5 mM EDTA, and 5 mM TCEP. PPARγ LBD was prepared at 50 μM in identical buffer. 100 μM Pio or PioOH were added to PPARγ LBD and TRAP220 and incubated on ice 30 minutes before each experiment. TRAP220 peptide (syringe) was titrated into PPARg LBD (sample cell). 20 total injections were made per experiment (0.4 μL for the first injection, 2.0 pL for subsequent injections), using a mixing speed of 1200 rpm, a reference power of 5 μcal/second, and a cell temperature of 25°C. Two runs were performed for each ligand-bound condition. Experiments were performed using a MicroCal iTC200 (Malvern). Data were processed in NITPIC (46) and analyzed by unbiased global fitting of both replicate runs per ligand-bound condition in SEDPHAT (47), followed by export to GUSSI for publication-quality figure preparation (48).

### NMR spectroscopy

Two dimensional [^1^H,^15^N] -transverse relaxation optimized spectroscopy (TROSY)-heteronuclear single quantum correlation (HSQC) data were collected at 298K using a Bruker 700 Mhz NMR instrument equipped with a QCI cryoprobe. Samples contained 200 μM ^15^N-labeled PPARγ LBD in a buffer (NMR buffer) containing 20 mM potassium phosphate (pH 7.4), 50 mM potassium chloride, 0.5 mM EDTA, 5 mM TCEP, and 10% D2O in the absence or presence of two molar equivalents of Pio or PioOH. Data were processed and analyzed using Topspin 3.0 (Bruker) and NMRViewJ (OneMoon Scientific, Inc.) (49). NMR analysis was performed using previously described rosiglitazone-bound NMR chemical shift assignments (BMRB entry 17975) for well resolved residues with consistent NMR peak positions via the minimum chemical shift procedure (11,28).

### X-ray crystallography

PPARγ LBD was incubated with 1-hydroxypioglitazone at a 1:3 protein/ligand molar ratio in PBS overnight before being concentrated to 10 mg/ml. Crystals were obtained after 7-10 days at 22°C by sitting-drop vapor diffusion against 50 μL of well solution. The crystallization drops contain 1 μL of protein sample mixed with 1μL of reservoir solution containing 0.1 M Tris-HCl, 0.8 M sodium citrate at pH 7.6. Crystals were flash-cooled in liquid nitrogen before data collection. Data collection was carried out at Beamline 5.0.2 at Berkeley Center for Structural Biology (Advanced Light Source). Data were processed, integrated, and scaled with the programs Mosflm and Scala in CCP4 (50,51). The structure was solved at 1.88À by molecular replacement using the program Phaser (52) that was implemented in the PHENIX package (53) using a previously published PPARγ LBD crystal structure (PDB code: 1PRG (54)) as the search model. The structure was refined using PHENIX with several cycles of interactive model rebuilding in COOT (55).

### Hydrogen-deuteirum exchange mass spectrometry (HDX-MS)

Solution-phase amide HDX experiments were carried out with a fully automated system described previously (56) with slight modifications. Five μl of PPARγ LBD protein (10 μM), without or with Pio or PioOH (100 μM), was mixed with 20 μL of D_2_O-containing NMR buffer and incubated at 4 °C for a range of time points (0s, 10s, 30s, 60s, 900s or 3,600s). Following exchange, unwanted forward or back exchange was minimized and the protein was denatured with a quench solution (5 M urea, 50 mM TCEP and 1% v/v TFA) at 1:1 ratio to protein. Samples were then passed through an in-house prepared immobilized pepsin column at 50 μL min^-1^ (0.1% v/v TFA, 15 °C) and the resulting peptides were trapped on a C_18_ trap column (Hypersil Gold, Thermo Fisher). The bound peptides were then gradient-eluted (5-50% CH3CN w/v and 0.3% w/v formic acid) across a 1 mm × 50 mm C_18_ HPLC column (Hypersil Gold, Thermo Fisher) for 5 min at 4 °C. The eluted peptides were then analyzed directly using a high resolution Orbitrap mass spectrometer (Q Exactive, Thermo Fisher). Each HDX experiment was performed in triplicate. To identify peptides, MS/MS experiments were performed with a Q Exactive mass spectrometer over a 70 min gradient. Product ion spectra were acquired in a data-dependent mode and the five most abundant ions were selected for the product ion analysis. The MS/MS *.raw data files were converted to *.mgf files and then submitted to Mascot (Matrix Science, London, UK) for peptide identification. Peptides with a Mascot score of 20 or greater were included in the peptide set used for HDX detection. The MS/MS Mascot search was also performed against a decoy (reverse) sequence and false positives were ruled out. The MS/MS spectra of all the peptide ions from the Mascot search were further manually inspected and only the unique charged ions with the highest Mascot score were used in estimating the sequence coverage. The intensity weighted average m/z value (centroid) of each peptide isotopic envelope was calculated with the latest version of our in-house developed software, HDX Workbench (57).

## FOOTNOTES

This work was supported by National Institutes of Health (NIH) grants R01DK101871 (DJK), R01DK105825 (PRG), and F32DK108442 (RB); National Science Foundation (NSF) funding to the Summer Undergraduate Research Fellows (SURF) program at Scripps Research [Grant 1659594]; and the Academic Year Research Internship for Undergraduates (AYRIU) program at Scripps Research.

